## Supplementary data for "Plasticity in centromere organization: Holocentromeres can consist of merely a few megabase-sized satellite arrays"

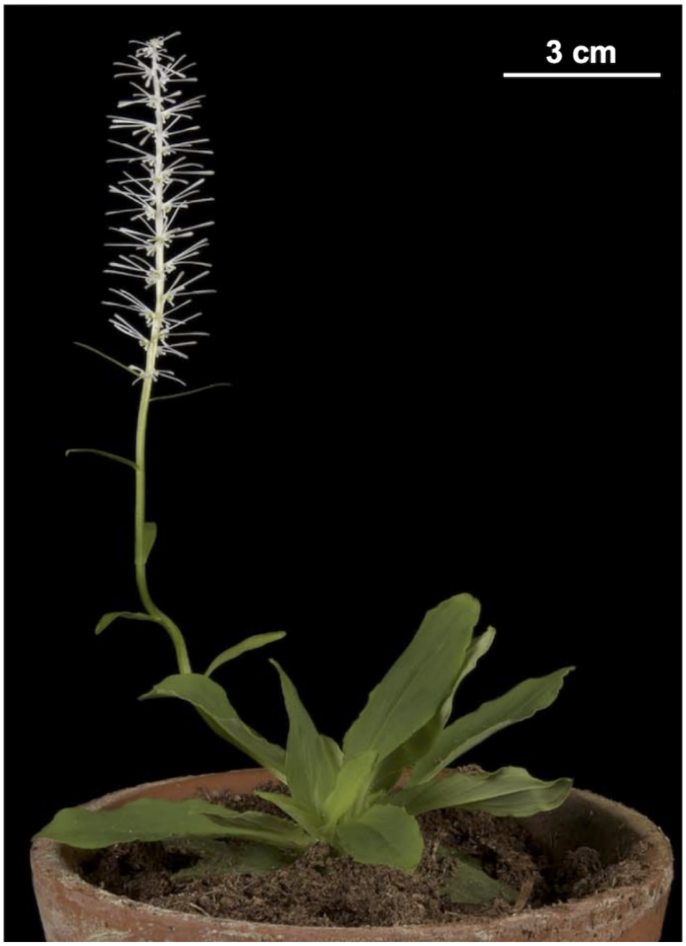


Supplementary Fig. 1

Flowering plant of *C. japonica.*

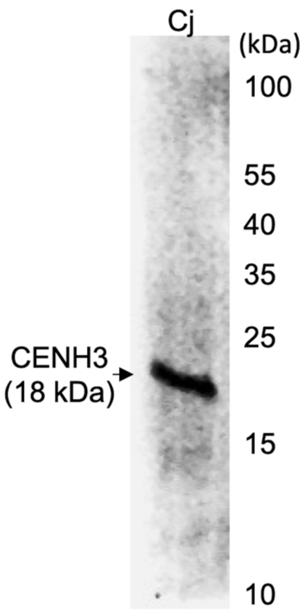


Supplementary Fig. 2

Western blot analysis of *C. japonica* CENH3. The specificity of the *C. japonica* anti-CENH3 antibody was confirmed by the detection of the predicted 18 kDa nuclear protein.

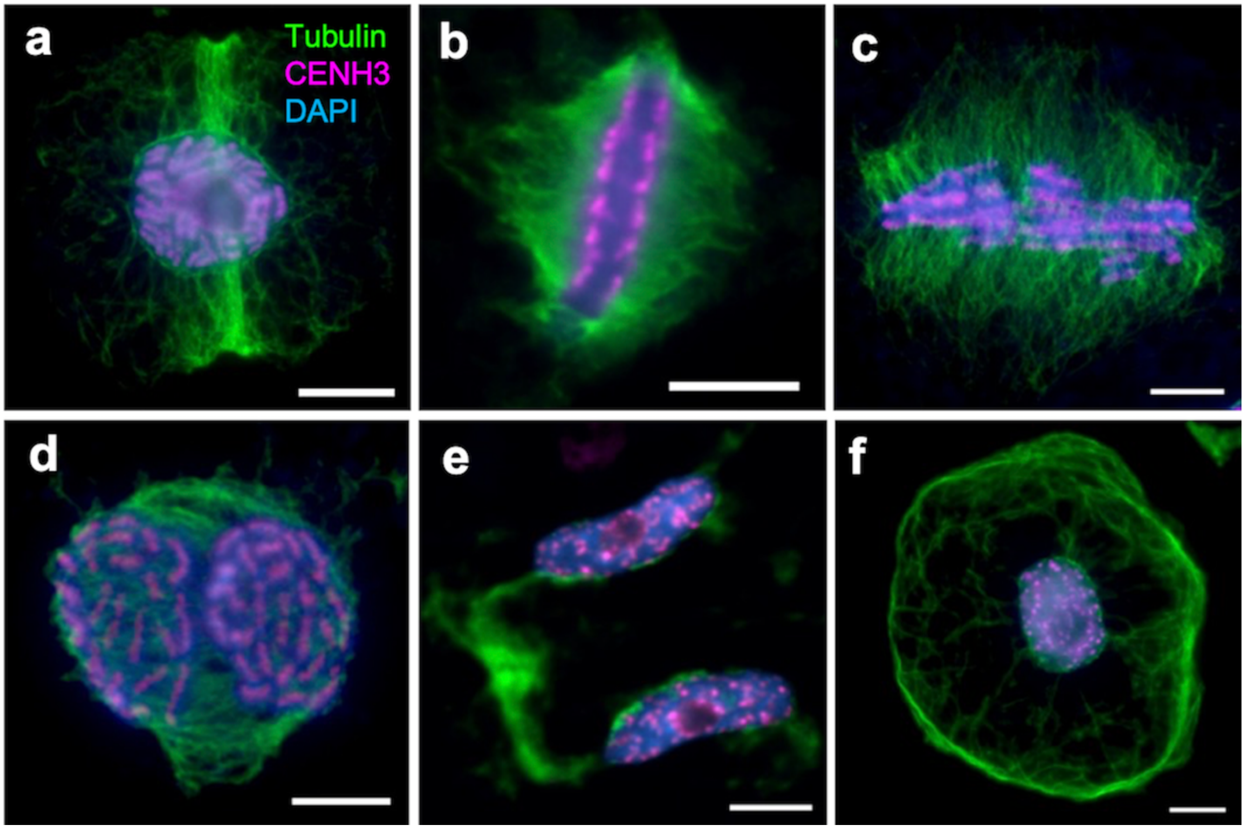


Supplementary Fig. 3

Immunolabelling of CENH3 (purple) and alpha-tubulin (green) in mitotic (a) prophase, (b) metaphase, (c) anaphase, (d) early telophase, (e) late telophase, and (f) interphase cells of *C. japonica.* Scale bar, 5 μm

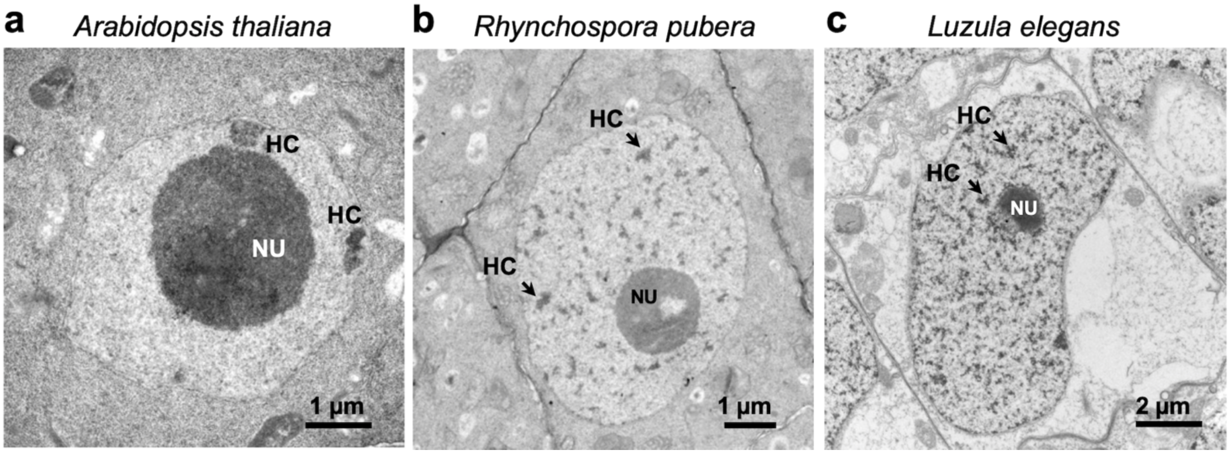


Supplementary Fig. 4

Transmission electron micrographs of root interphase nuclei of monocentric (a) *A. thaliana* and holocentric (b) *R. pubera* and (c) *L. elegans*. HC: heterochromatic chromocenter (arrows), NU: nucleolus

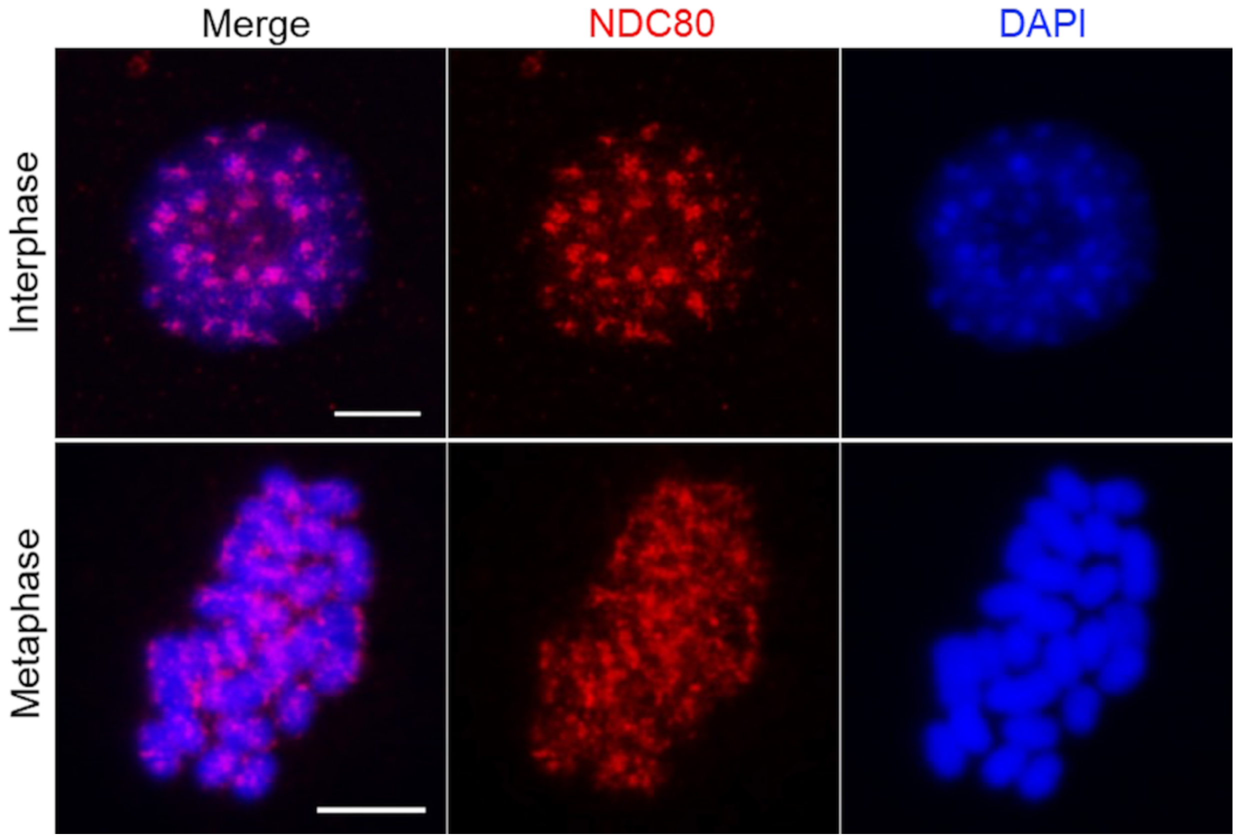


Supplementary Fig. 5

Immunodetection of the kinetochore protein NDC80 (red) in interphase nucleus and mitotic metaphase chromosomes of *C. japonica*. Nucleus and chromosomes were counterstained with DAPI and pseudocolored in blue. Scale bar, 5 μm

H2AThr120

```
Homo_sapiens_NP_003500             HLQLAIRNDEELNKLLGKVTIAQGGVLPNIQAVLLPKKTESHHKAKGK----------------
Arabidopsis_thaliana_NP_198119     HLCLAIRNDEELGRLLHGVTIASGGVLPNINPVLLPKKSTASSSQAEK------ASATKSPKKA
Zea_mays_H2A_P40280                HVLLAIRNDEELGKLLGGVTIAHGGVLPNINPVLLPKKTAEKASSGGSKEAKSPKKAAKSPKKA
Chionographis_japonica_H2A_6247    HVLLAVRNDEELGKLLAGVTIAHGGVLPNINPVLLPKKASN-----SGKEPKSPTKATKSPKKA
Chionographis_japonica_H2A_6244    HVLLAVRNDEELGKLLAGVTIAHGGVLPNINPVLLPKKASN-----SGKEPKSPTKATKSPKKP
Phoenix_dactylifera_H2A_XP_026656918 HVLLAVRNDEELGKLLAGVTIAHGGVLPNINPVLLPKKTA-------SKEPKSPSKATKSPKKA
Chionographis_japonica_H2A_6251    HVLLAVRNDEELGKLLSGVTIAHGGVLPNINPVLLPKKSA------SAKEPKSPSKATKSPKKA
```

Supplementary Fig. 6

Sequence alignment of histone H2A of different species. The highly conserved threonine 120 (Thr120) of H2A sequence in *C. japonica* is changed to either alanine (A) or serine (S). The translated sequence of three H2A transcripts identified from the *C. japonica* root transcriptomes were used in the alignment.

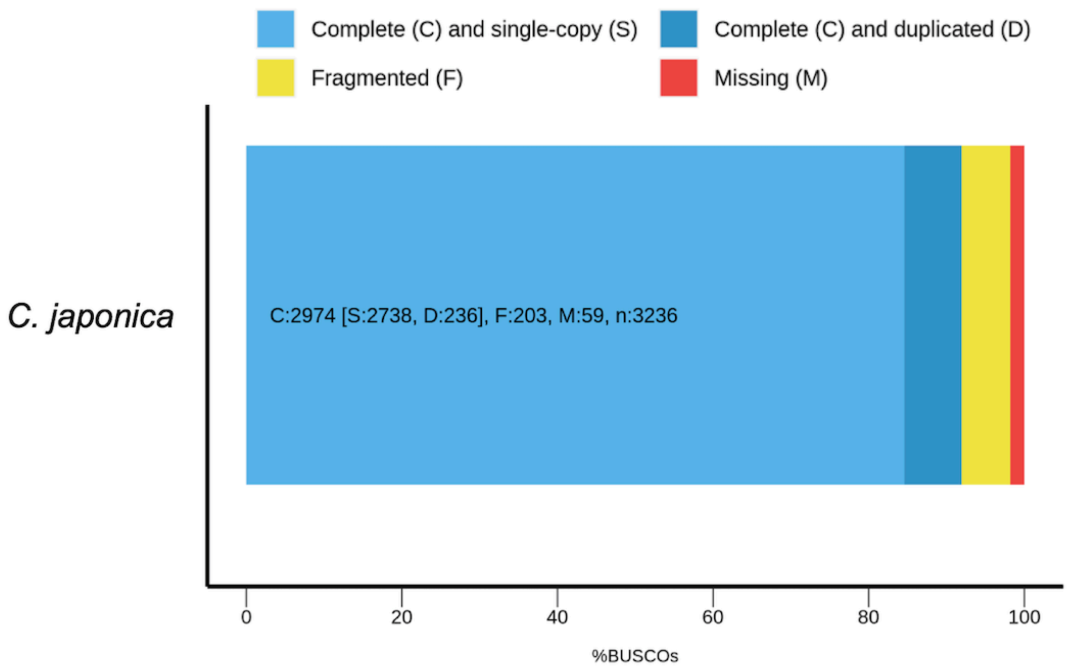


Supplementary Fig. 7

BUSCO assessment of the *C. japonica* assembled genome. The completeness of the assembled genome was assessed using BUSCO analysis against the Liliopsida_odb10 dataset.

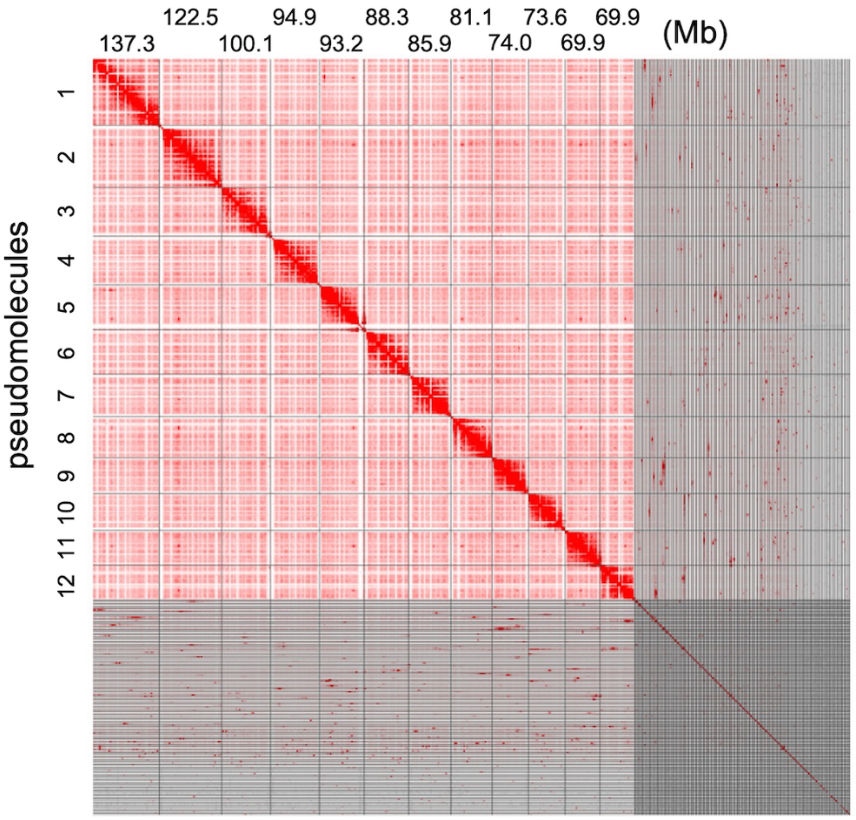


Supplementary Fig. 8

Hi-C map for the assembled pseudomolecules and contigs of *C. japonica*. The size of the 12 pseudomolecules is indicated.

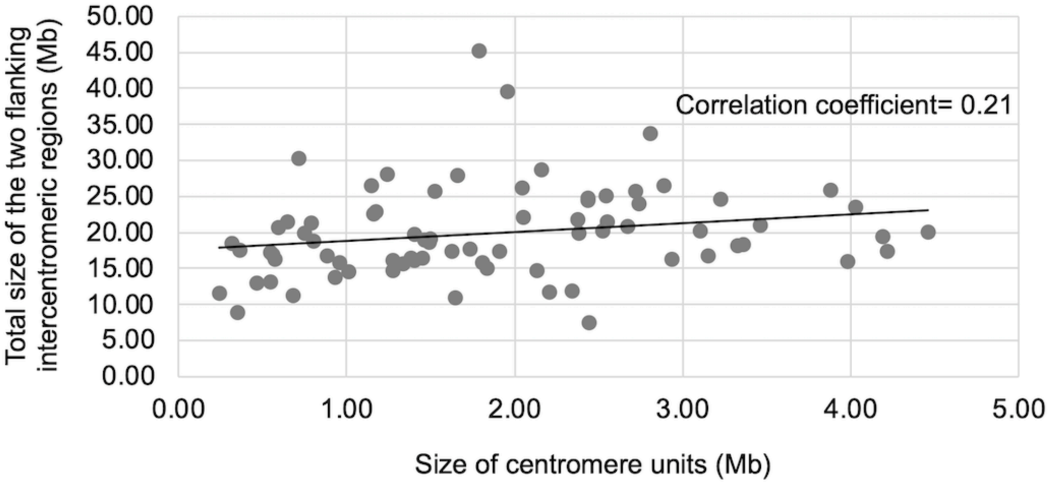


Supplementary Fig. 9

Assessment of the correlation between the size of centromere units and two flanking intercentromeric regions. A low correlation was revealed by the correlation coefficient of 0.21.

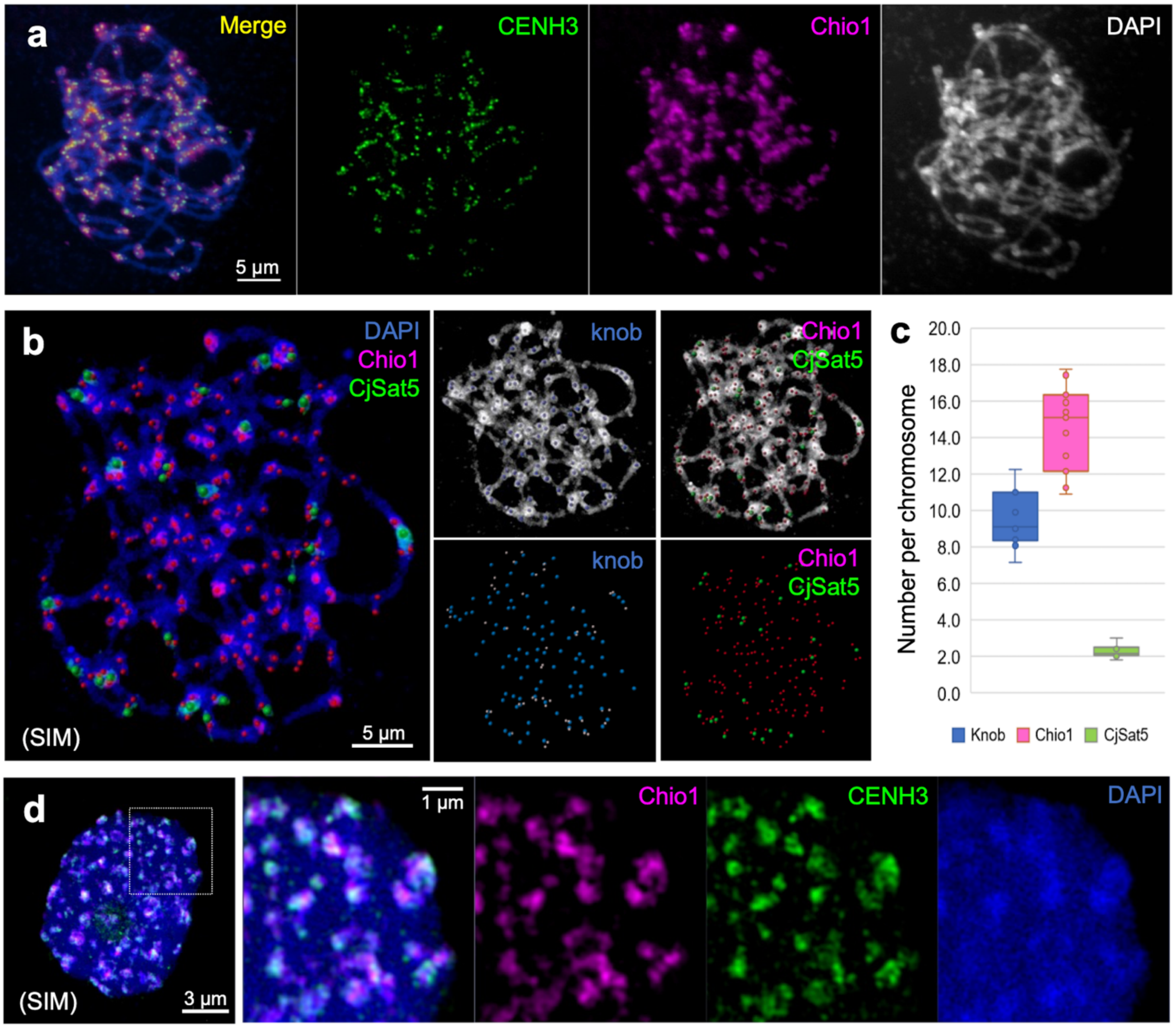


Supplementary Fig. 10

The distribution of centromere units in *C. japonica*. (a) Immuno-FISH shows colocalization of the centromeric CENH3 (green) and Chio1 satellite repeat (purple) in pachytene chromosomes. (b) The knob structure (blue), signals of Chio1 (purple) and CjSat5 (green) satellite repeats were detected using the 'Spots' tool of Imaris 9.7 (Oxford instruments, UK), and (c) the average number of knobs, Chio1 and CjSat5 signals per chromosome were calculated in 10 pachytene spreads. (d) The colocalization of CENH3 (green) and Chio1 (purple) in an interphase nucleus. The percentage of overlapped signals was ~65%, from 56.5% to 74.4% (n=11) measured by the Coloc tool of Imaris. Pachytene chromosomes and interphase nucleus were counterstained with DAPI.

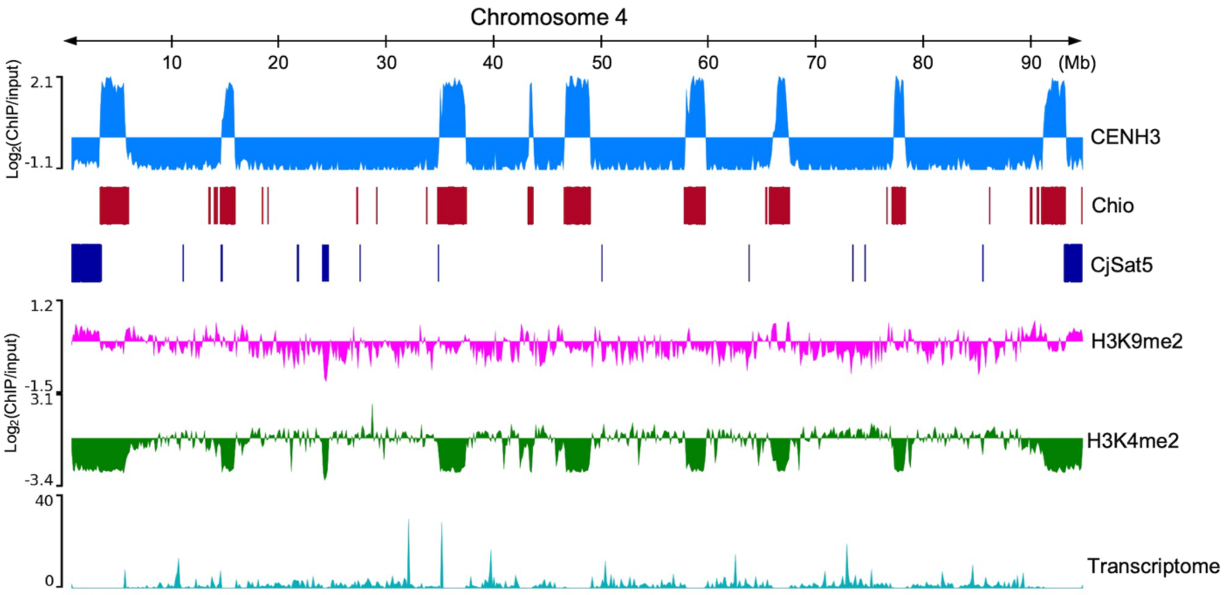


Supplementary Fig. 11

The enrichment of CENH3-, H3K9me2-, and H3K4me2-ChIPseq and distribution of the centromeric Chio satellite arrays, subtelomeric CjSat5 satellite repeat, and root RNAseqs. The ChIPseq signal tracks are represented as the average of $\log_2$ ratio of ChIP/input in genome-wide 1 kb windows. Root transcriptome signal is shown as normalized read per kilobase per million (RPKM) in 1 kb windows.

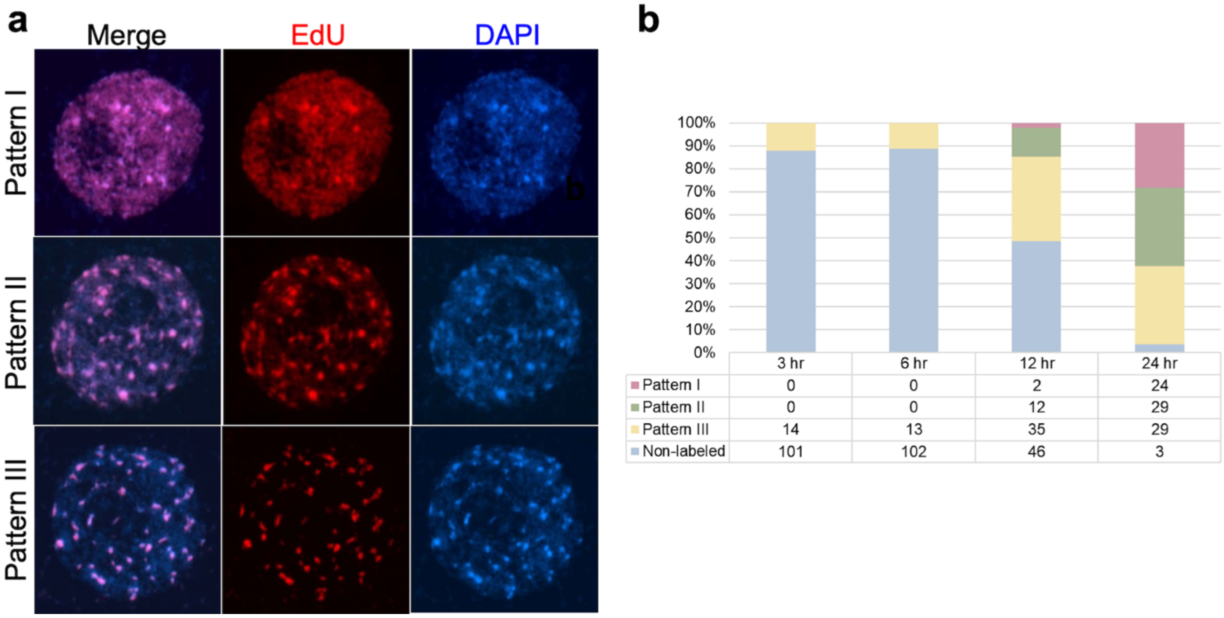


| | 3 hr | 6 hr | 12 hr | 24 hr |
|---|---|---|---|---|
| Pattem I | 0 | 0 | 2 | 24 |
| Pattem II | 0 | 0 | 12 | 29 |
| Pattem III | 14 | 13 | 35 | 29 |
| Non-labeled | 101 | 102 | 46 | 3 |

Supplementary Fig. 12

EdU labeling-based DNA replication analysis of *C. japonica*. (a) The interphase replicating pattern I to III correspond to the early, mid, and late S phase, respectively. (b) The number of each pattern in the fixed materials with pulse recovery times of 3, 6, 12, and 24 hours counted in metaphase spreads are listed. The bar plot shows the percentage of each pattern in different samples and number of counted metaphase chromosome spreads is indicated in the table.

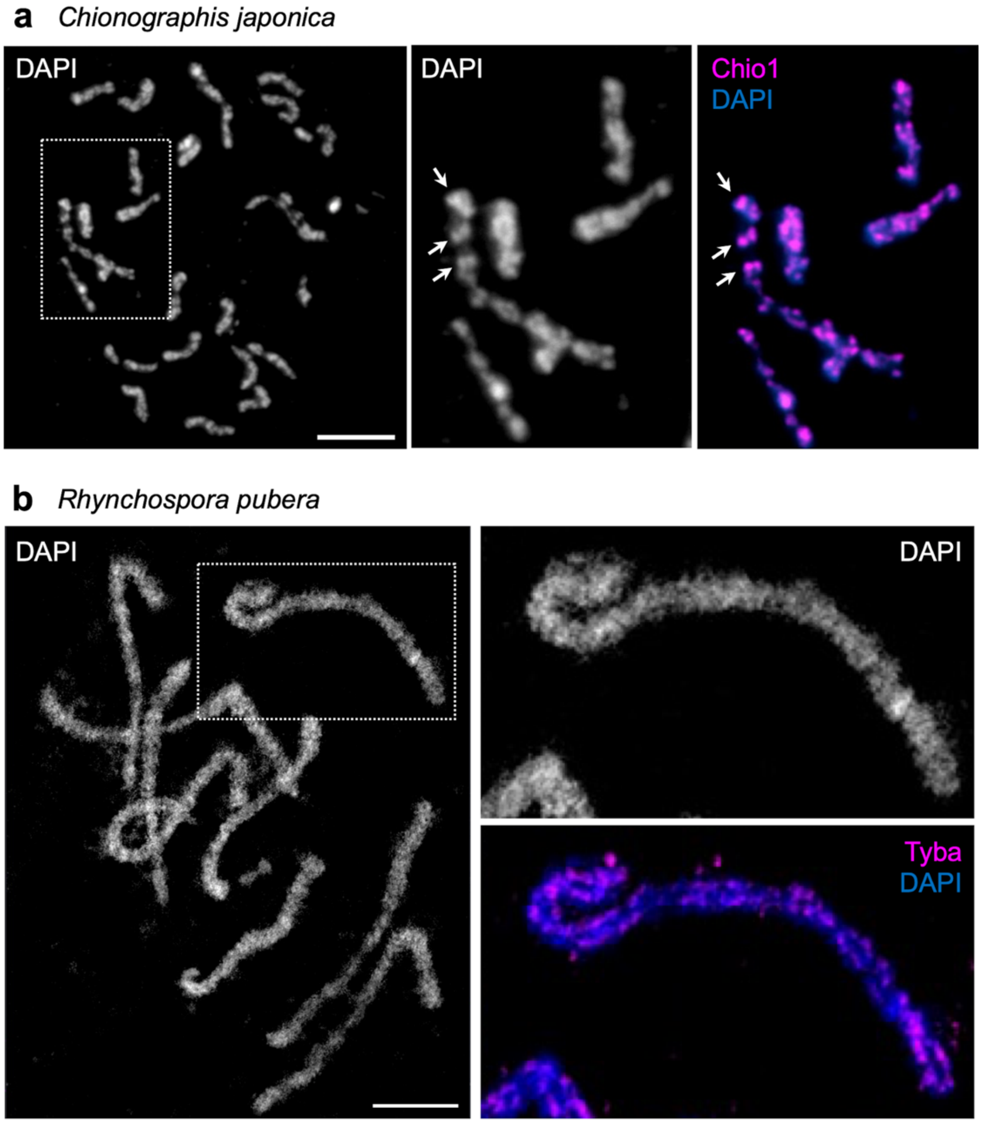


Supplementary Fig. 13

Chromosome morphology of *C. japonica* and *R. pubera* during mitotic condensation. (a) The prometaphase chromosomes of *C. japonica* are non-uniformly condensed. Centromeric Chio1 repeats (purple) cluster and colocalize with heterochromatic regions. Enlargements (squares) are shown in the right panels. (b) In contrast, the prometaphase chromosomes of *R. pubera* show a uniform structure and line-like holocentromere-specific Tyba signals. Chromosomes were counterstained with DAPI. Scale bar, 5 µm

Supplementary Table 1

Number of CENH3 signal clusters in interphase nuclei of *C. japonica* counted in 2D and 3D stacked images

| Root G1 nuclei* | No. of CENH3 signal clusters per nucleus | No. of CENH3 signal clusters per chromosomes |
|---|---|---|
| 2D image-1 | 64 | 2.67 |
| 2D image-2 | 49 | 2.04 |
| 2D image-3 | 74 | 3.08 |
| 2D image-4 | 82 | 3.42 |
| 2D image-5 | 79 | 3.29 |
| 2D image-6 | 72 | 3.00 |
| 2D image-7 | 57 | 2.38 |
| 2D image-8 | 59 | 2.46 |
| 2D image-9 | 59 | 2.46 |
| 2D image-10 | 101 | 4.21 |
| 2D image-11 | 70 | 2.92 |
| 2D image-12 | 50 | 2.08 |
| 2D image-13 | 107 | 4.46 |
| 2D image-14 | 76 | 3.17 |
| 2D image-15 | 76 | 3.17 |
| 2D image-16 | 94 | 3.92 |
| 2D image-17 | 68 | 2.83 |
| 2D image-18 | 62 | 2.58 |
| 2D image-19 | 80 | 3.33 |
| 2D image-20 | 58 | 2.42 |
| 2D image-21 | 71 | 2.96 |
| 2D image-22 | 58 | 2.42 |
| 2D image-23 | 79 | 3.29 |
| 2D image-24 | 67 | 2.79 |
| 2D image-25 | 70 | 2.92 |
| 2D image-26 | 49 | 2.04 |
| 2D image-27 | 48 | 2.00 |
| 2D image-28 | 55 | 2.29 |
| 2D image-29 | 63 | 2.63 |
| 2D image-30 | 48 | 2.00 |
| **Average** | **68.17** | **2.85** |
| 3D image-1 | 156 | 6.50 |
| 3D image-2 | 55 | 2.29 |
| 3D image-3 | 47 | 1.96 |
| 3D image-4 | 115 | 4.79 |
| 3D image-5 | 66 | 2.75 |
| 3D image-6 | 51 | 2.13 |
| 3D image-7 | 43 | 1.79 |
| 3D image-8 | 82 | 3.42 |
| 3D image-9 | 62 | 2.58 |
| 3D image-10 | 48 | 2.00 |
| 3D image-11 | 43 | 1.79 |
| 3D image-12 | 41 | 1.71 |
| **Average** | **67.42** | **2.81** |

* The G1 nuclei were isolated from roots of *C. japonica,* followed by sorting using flow cytometry

Supplementary Table 2

Summary of the *de novo* genome assembly of *C. japonica*

| | *Chionographis japonica* | |
|---|---|---|
| Genome size (Mb/1C)* | 1,368 | |
| GC content (%) | 41.26 | |
| Sequence coverage | 58.5× | |
| Scaffolding strategy | Hi-C | |
| | Assembly | Scaffolding |
| Number of contigs/ scaffolds | 3,786 | 3,263 |
| Assembly size (bp) | 1,526,137,861 | 1,526,120,517 |
| Longest contig (bp) | 11,690,282 | 137,288,582 |
| N50 (bp) | 2,877,649 | 81,106,678 |
| N75 (bp) | 1,067,322 | 2,104,563 |
| L50 | 150 | 8 |
| L75 | 373 | 32 |

*The genome size was determined by flow cytometry

Supplementary Table 3

The size and centromere characterization on the 12 chromosome scaffolds of *C. japonica.*

| Chromosome scaffold | Total length (Mb) | No. of centromere units | Average size of centromere units (Mb) | Average interval between centromere units (Mb) |
|---|---|---|---|---|
| Chr 1 | 137.29 | 11 | 2.09 | 11.20 |
| Chr 2 | 122.50 | 10 | 1.65 | 11.59 |
| Chr 3 | 100.11 | 8 | 2.29 | 11.35 |
| Chr 4 | 94.91 | 9 | 1.76 | 9.34 |
| Chr 5 | 93.21 | 11 | 1.46 | 7.58 |
| Chr 6 | 88.32 | 7 | 2.40 | 11.64 |
| Chr 7 | 85.88 | 8 | 2.30 | 9.25 |
| Chr 8 | 81.11 | 7 | 2.38 | 10.74 |
| Chr 9 | 74.04 | 8 | 1.40 | 8.98 |
| Chr 10 | 73.57 | 7 | 1.78 | 8.59 |
| Chr 11 | 69.90 | 7 | 1.27 | 10.17 |
| Chr 12 | 69.89 | 7 | 1.93 | 9.18 |
| Average | 90.89 | 8.3 | 1.89 | 9.97 |
| Sum | 1090.73 | 100 | - | - |

Supplementary Table 4

Characterization of high-copy satellite repeats of *C. japonica* used for FISH.

| Satellite repeat | Monomer (bp) | Genome proportion (%) | Sequence of oligo probe/ PCR primer (5'-3') |
|---|---|---|---|
| Chio1 | 23 | 16.11 | TCATTCGTACGATCCATTCTAAT |
| Chio2 | 28 | | TCATTCGTACGAATGTAGCCATTCTAAT |
| CjSat3 | 152 | 0.59 | F: ACACCCTCTAAGAGCCTCGC<br>R: AGCCAAAACCGCTCCATATT |
| CjSat4 | 181 | 0.88 | F: CGGACTTGCGAGCGAGTT<br>R: ATGCCGATCCGATGACGAT |
| CjSat5 | 270 | 2.20 | F: CTCGACATGTTCGTGCTGAT<br>R: CCAGTCACAAGAAAACGGAGA |

Supplementary Table 5

Protocol for combined conventional and microwave-assisted fixation, dehydration and embedding in Spurr resin of root tips and leaf cuttings suitable for TEM.

| **Combined conventional & microwave assisted root tissue preparation in a PELCO Bio Wave®34700-230 (Ted Pella, Inc., Redding CA, USA)** | | | | |
|---|---|---|---|---|
| **Process** | **Reagent** | **Power [W]** | **Time [sec]** | **Vacuum [mm Hg]** |
| **1. Primary Fixation** | 2.0% (v/v) glutaraldehyde and 2.0% (v/v) paraformaldehyde in 0.05 M cacodylate buffer (pH 7.3) | 150<br>0<br>150<br>0<br>150 | 60<br>60<br>60<br>60<br>60 | 0<br>0<br>0<br>0<br>0 |
| **2. Wash** | 1x 0.05 M cacodylate buffer (pH 7.3) and 2x aqua dest. | 150 | 60 | 0 |
| **3. Secondary fixation** | 1% (v/v) osmiumtetroxide in aqua dest. | 0<br>80<br>0<br>80 | 60<br>120<br>60<br>120 | 10<br>10<br>10<br>10 |
| | samples were kept for additional 15 min on a shaker | | | |
| **4. Wash** | 3x aqua dest. | 150 | 60 | 0 |
| **5. Dehydration** | acetone: 30%, 40%, 50%, 60%, 70%, 80%, 90%, 2x 100% and 1 x propylenoxide. | 150 | 60 | 0 |
| | after each step samples were kept for additional 2 min on a shaker | | | |
| **6. Resin Infiltration** | 25% Spurr resin in propylenoxide<br>50% Spurr resin in propylenoxide<br>75% Spurr resin in propylenoxide<br>100% Spurr resin | overnight on shaker at RT<br>4 hrs on shaker at RT<br>4 hrs on shaker at RT<br>overnight on shaker at RT | | |
| **7. Polymerisation** | 24 hrs at 70°C in prepolymerized flat embedding moulds in a heating cabinet | | | |
